# Estimating the Preferred Walking Speeds Based on the Center of Mass Work Analysis

**DOI:** 10.1101/2024.07.16.603701

**Authors:** Seyed-Saleh Hosseini-Yazdi, John EA Bertram

## Abstract

This study investigates the role of optimal push-off impulses in minimizing the total mechanical work dissipation per step, aiming to achieve passive single support work performance for a variety of walking conditions. By optimizing push-offs to cover the entire step’s energy demands, the single support work may become only storage and release of mechanical energy through tendons and tissues. Our simulations indicate that for each walking speed, there is an optimal push-off impulse that ensures energy balance without the need for additional energy input or dissipation during the subsequent single support phase when the soft tissue dissipation is negligible. For this circumstance, the step total dissipation is minimum. For level ground (even terrain), the estimated preferred walking speeds align closely with literature values for young adult (∼ 1.2 m.s^−1^)and older adult (∼ 1.0 m.s^−1^) self-selected speeds, with a reduction observed for the restricted view of oncoming irregularities in the substrate terrain (15%). For walking on uneven surfaces, terrain amplitude was shown to impact walking costs quadratically, with optimal speeds declining by approximately 20% per unit increase in terrain amplitude. Key findings include evidence that net single support work remains near zero when push-off optimally covers collision and gravity work, confirming the passive nature of single support work under this condition. We also observed that the preferred speeds for older adults tend to be 12-15% lower than for younger adults, likely due to biomechanical adaptations. Beyond certain terrain amplitudes, no preferred walking speed allowed fully passive single support work, highlighting a possible biomechanical threshold where ankle push-off alone becomes insufficient and hip torque compensation may be necessary. This approach provides a framework for estimating preferred walking speeds across different terrain amplitudes (continuous parameter), varying conditions and demographics, with potential applications in designing assistive devices and gait rehabilitation protocols that reduce metabolic cost through optimal mechanical work management.

## Introduction

One primary objective of walking is to minimize energy expenditure [1] while simultaneously achieving other task-specific goals [2]. Research suggests that energy expenditure is minimized when humans walk at a nominal or preferred walking speed (self-selected) [3]. The preferred walking speed is traditionally defined as the speed at which the cost of transport is minimized [4]; walking faster or slower than this speed results in increased energetic costs [5]. To determine an individual’s average preferred walking speed based on energetics, subjects walk at various velocities while their energy expenditure is measured [6]. The walking speed that coincides with the minimum of the resulting parabolic cost of transport trajectory is identified as the preferred walking speed [2], [4]. In over ground free walking, the self-selected speed is naturally adopted. This speed must reflect an optimal balance between energy expenditure and comfort for the individual [3]. Since it has been proposed that the active center of mass (COM) work during step-to-step transitions closely represents metabolic walking cost [7], a mechanical approach may also estimate preferred walking speed based on COM mechanical work.

It is hypothesized that walking economy is optimized when an appropriate pre-emptive push-off energizes gait progression [8]. The push-off redirects the COM velocity from the previous step upward, allowing the subsequent collision to adjust the COM velocity to match the new stance [8]. In a properly timed pre-emptive push-off, the collision loss is minimized and is equal to the push-off work energy infusion (Figure 1A) [9], [10]. When the push-off is suboptimal, it only partially energizes the gait, leading to a greater collision dissipation at mid-transition to adjust the COM trajectory for the next step (Figure 1B). This leads to a reduced outcome velocity after the transition [9], requiring additional positive mechanical work be added to sustain the gait. Otherwise, the COM speed will decline, and the gait may become unsustainable. This extra compensatory work likely occurs during the single support phase, where the added positive work is termed ‘rebound’ (Figure 1D) [10], [11]. Similarly, if push-off exceeds the subsequent heel-strike collision, the post-transition velocity increases. Thus, to maintain the average velocity, energy dissipation during the subsequent step is required. This dissipation must occur when the single support negative work occurs (preload, Figure 1D).

**Figure 1.**
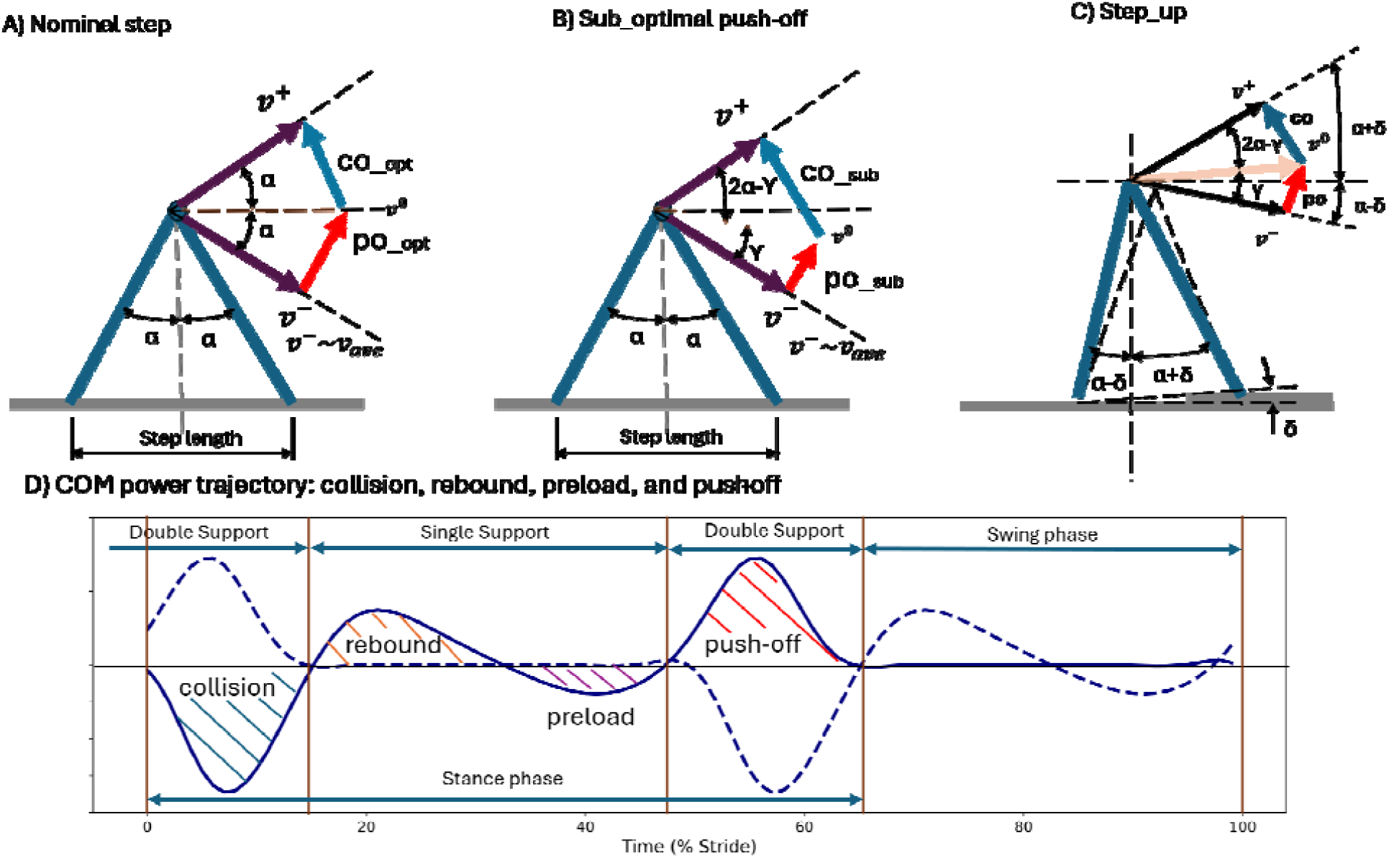
The step-to-step transition is when the stance leg changes during double stance. (A) During nominal walking, the push-off work is exerted pre-emptively and is equal to the subsequent collision. (B) If a sub-optimal push-off is exerted, the subsequent collision magnitude grows. The reverse is true when the push-off is more than the nominal magnitude (C) Uneven walking is a deviation from nominal even walking. In the ideal case, the pre-emptive push-off must provide the mechanical energy for step change and gravity work. (D) The human walking COM power trajectory depicts four distinct regions: collision (heel-strike), rebound, preload, and push-off (toe-off). Since both limbs perform work on the COM during the double stance, the other leg’s power trajectory is demonstrated with a dashed line. The negative work components (collision and preload) and positive work components (rebound and push-off) determine the work balance during contact, influencing the energetic efficiency of the gait and the compensations required.

In nominal walking, it is proposed that single support work is passive, occurring through mechanical energy storage and release facilitated by tendons and other tissues [10], [12]. Thus, the positive energy generated by the rebound phase must be absorbed during the preload (Figure 1D) [10]. During the single support phase (rigid body), when the COM motion resembles an inverted pendulum [13], the total work contributions of the leg joints have been shown to balance rebound and preload in nominal walking with minimal soft tissue dissipation [14]. To counteract deficits from push-off and collision [11], if the rebound is to energize the COM progression, it must exceed the preload; conversely, if dissipation is required, preload must exceed rebound. Hence, it is plausible that even during normal level walking, certain steps may deviate from the preferred regime, causing push-off, collision, rebound, or preload to become unequal.

Walking on an uneven substrate (uneven walking) presents a case where push-off may deviate from the optimal pattern (Figure 1C). For example, heel strike may occur earlier than expected [15], particularly in the absence of visual cues about upcoming terrain variations [16]. In such cases, the collision occurs before the prior push-off fully adjusts the COM velocity [10], [17], increasing the collision dissipation magnitude and demanding additional active energy input [16], likely through hip torque [9], [18]. Age and a restrictive view of upcoming surface changes (here referred to as lookahead) also influences walking biomechanics [16], [19]. Thus, we propose that for a given walking condition, there is a walking speed at which COM mechanical work dissipation is optimized. Based on a powered simple walking model [9], we propose that, in the mechanically optimal speed, push-off that requires active muscle work, provides all necessary walking energy. As such, no energy adjustment (compensation or dissipation) is required after the step transition. Consequently, rebound serves only to straighten the stance leg and swing the limbs [10]. It is fully absorbed by the negative preload work and remains passive [10], [12].

Since during uneven walking the COM elevation change for each step is unknown, it is challenging to estimate the optimal push-off as the sum of step dissipation and gravitational work. We therefore propose “rebound work equal to preload”—where single support work is suggested to be passive—as a criterion for identifying the mechanically preferred walking speed.

## Materials and Methods

### Simulation

Building on a powered simple walking model [9], we compute the analytical collision dissipation when push-off is suboptimal (a fraction of the nominal push-off). As shown in Figure 1B, the velocity prior to transition is approximated by the average velocity (*v*_*ave*_) and ‘2_α_’ represents the leading and trailing leg angles at the point of transition [9]. The suboptimal push-off (*po*_*sub*_) modifies the pre-transition COM velocity (*v*_*ave*_) partially by the angle of ‘*γ*’ which is less than ‘*α*’:

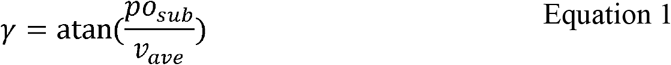

With the mid-transition COM velocity denoted as *v*^0^, we can estimate the collision impulse (*co*) as:

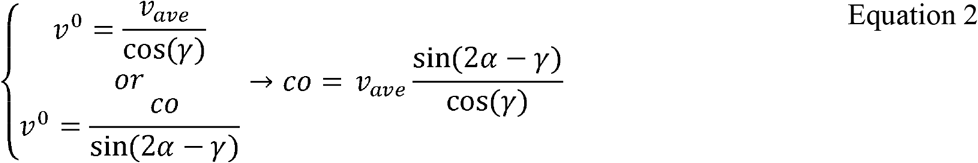

For uneven walking, when the walker steps up to a height Δ*h*, the post-transition velocity 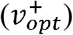 must exceed the pre-transition velocity to account for the increased potential energy. Thus:

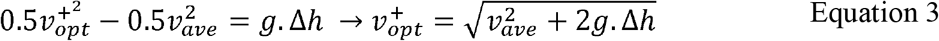

The mechanical work associated with a step transition impulse (*u*) is calculated as 0.5*u*^2^ [15]. In the optimal scenario, where all transition work is provided by a pre-emptive push-off, the transition impulses are:

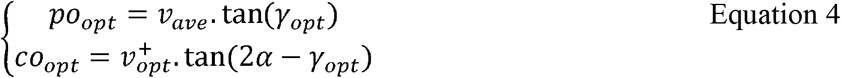

If PO and CO denote push-off and collision work, respectively, their difference should match the gravitational work needed to maintain momentum:

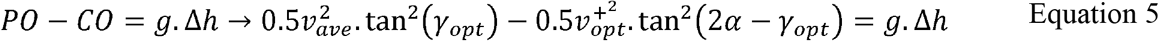

We solve Equation 6 numerically to determine *γ*_*opt*_ for given conditions. For our calculations, we assume *v*_*ave*_ = 1.25 m.s^−1^, and its corresponding α =0.35 [12]. For uneven walking, Δ*h* = 0.025m is used. We ran simulations for even and uneven walking where the push-off impulse varied from zero to the optimal level (Figure 2). Results were normalized by the optimal push-off work for each case.

**Figure 2.**
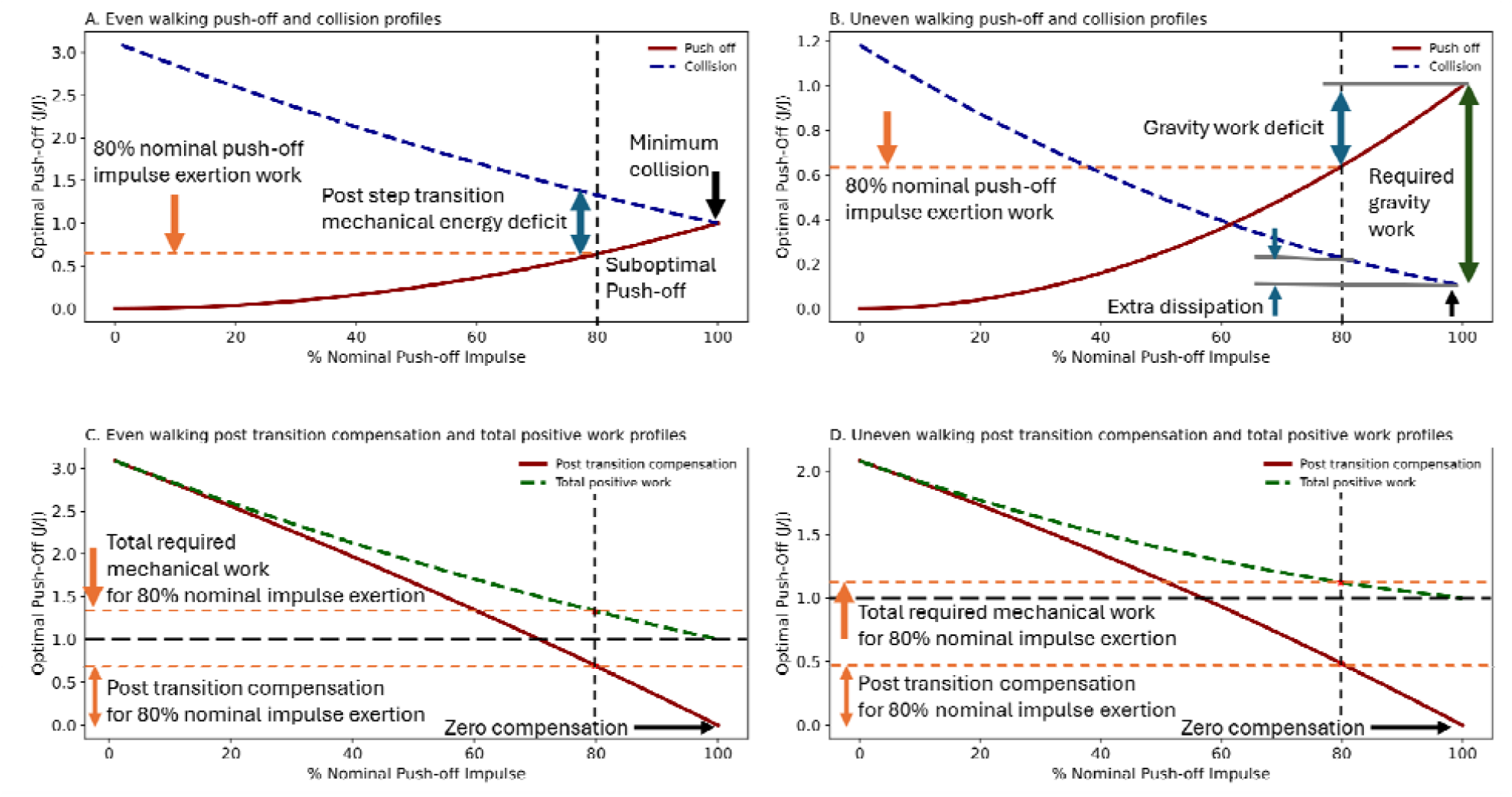
Simulation results when suboptimal push-off is exerted: (top row) resulting collision work trajectories during the even walking and with a known step up (uneven walking with known gravity work). The exerted push-off impulse ranges from zero to nominal magnitudes. (Bottom row) post transition work compensation and the total positive work trajectories of suboptimal push-off impulse for even walking and known step up (uneven walking).

The simulation shows that suboptimal push-off leads to an increase in collision dissipation, creating a mechanical energy deficit post-transition. This deficit requires additional positive work during the subsequent single support phase [11]. For even walking, if only 80% of optimal push-off impulse is exerted, after transition, an additional 70% of nominal push-off work must be performed, increasing total positive work to 133% of nominal. This implies 33% extra mechanical work, with a corresponding metabolic cost.

For Δ*h* = 0.025m, the required gravitational work is approximately 90% of optimal push-off (magnitude of dark green arrow, Figure 2B). With only 80% of required push-off impulse, the post-transition mechanical energy compensation would be 49% of what’s needed to reach the same speed at the subsequent midstance (optimal push-off work). The resulting total work performance is 112% of optimal work. This indicates a need for 12% additional positive work with associated metabolic consequences. The simulation suggests increased rebound work beyond what preload can absorb when pre-emptive push-off fails to cover total transition energy. Thus, when PO – CO < 0 (even walking) or PO – CO < gravity work, the push-off is suboptimal.

Conversely, when the push-off exceeds the nominal value, the post-transition speed surpasses the average speed (speed at midstance, Figure 3), requiring dissipation to maintain steady speed. For even walking, when push-off impulse is 125% of nominal, a mechanical energy equal to 91% of nominal push-off work must be dissipated. Similar excess push-off raises the required dissipation to 65% of optimal work for uneven walking. Notably, the simple powered walking model defines a maximum pre-emptive push-off as: *po*_*max*_ = *v*_*ave*_ .tan(2*α*) [9].

**Figure 3.**
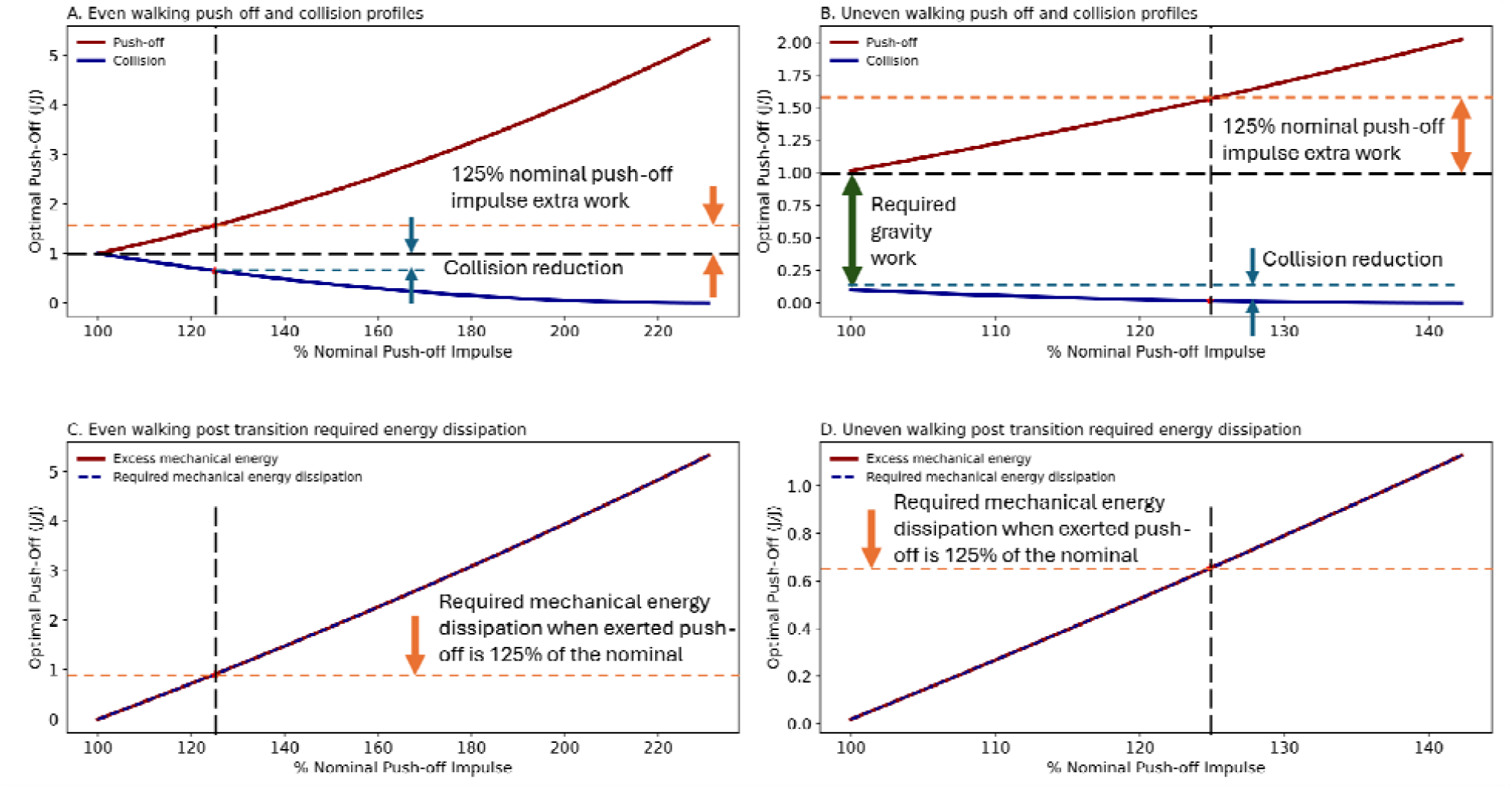
Simulation results for excess pre-emptive push-off: (top row) resulting collision work trajectories during even walking and known step up (uneven walking) as push-off impulse ranges from nominal magnitude to maximum analytical level (). For maximum analytical push-off impulse, the collision work becomes zero. (Bottom row) post transition requires work dissipation of the excess mechanical energy arising from the surplus push-off impulse for even walking and step up (uneven walking).

Therefore, the simulation demonstrates that for any given walking speed, suboptimal or excessive push-off, increases the magnitude of mechanical energy dissipation, leading to elevated positive mechanical work requirements (Figure 4). Since mechanical energy largely accounts for walking energetics [7], and energy conservation is a fundamental walking goal [1], it can be inferred that suboptimal or excessive push-off is not favorable from an energy perspective.

**Figure 4.**
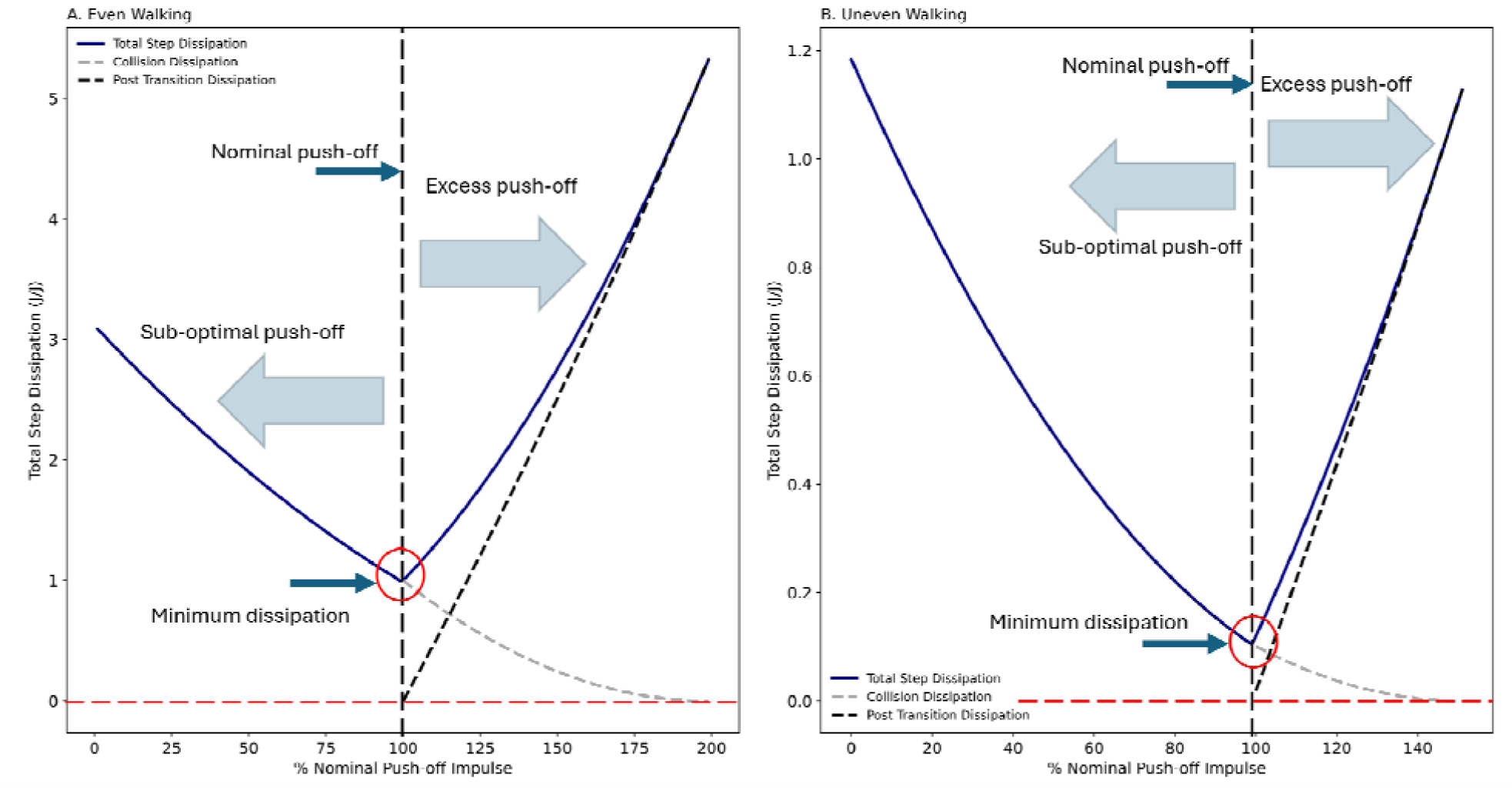
The total analytical dissipation, based on the magnitude of the exerted push-off impulse, is calculated for: (A) even walking and (B) uneven walking. For suboptimal push-off, the total dissipation is equal to the amplified collision. In contrast, for excessive push-off, the total dissipation comprises both step collision losses and the excess push-off work. The minimum dissipation occurs at the nominal push-off.

### COM Work Component Traces Based on Experimental Data

Using the trajectories proposed for various walking conditions—including state of lookahead, terrain amplitudes, and walking speeds—for both young and older adults [11], [20], we developed COM work trajectories (Figure 5 and Table 1).

**Table 1:**
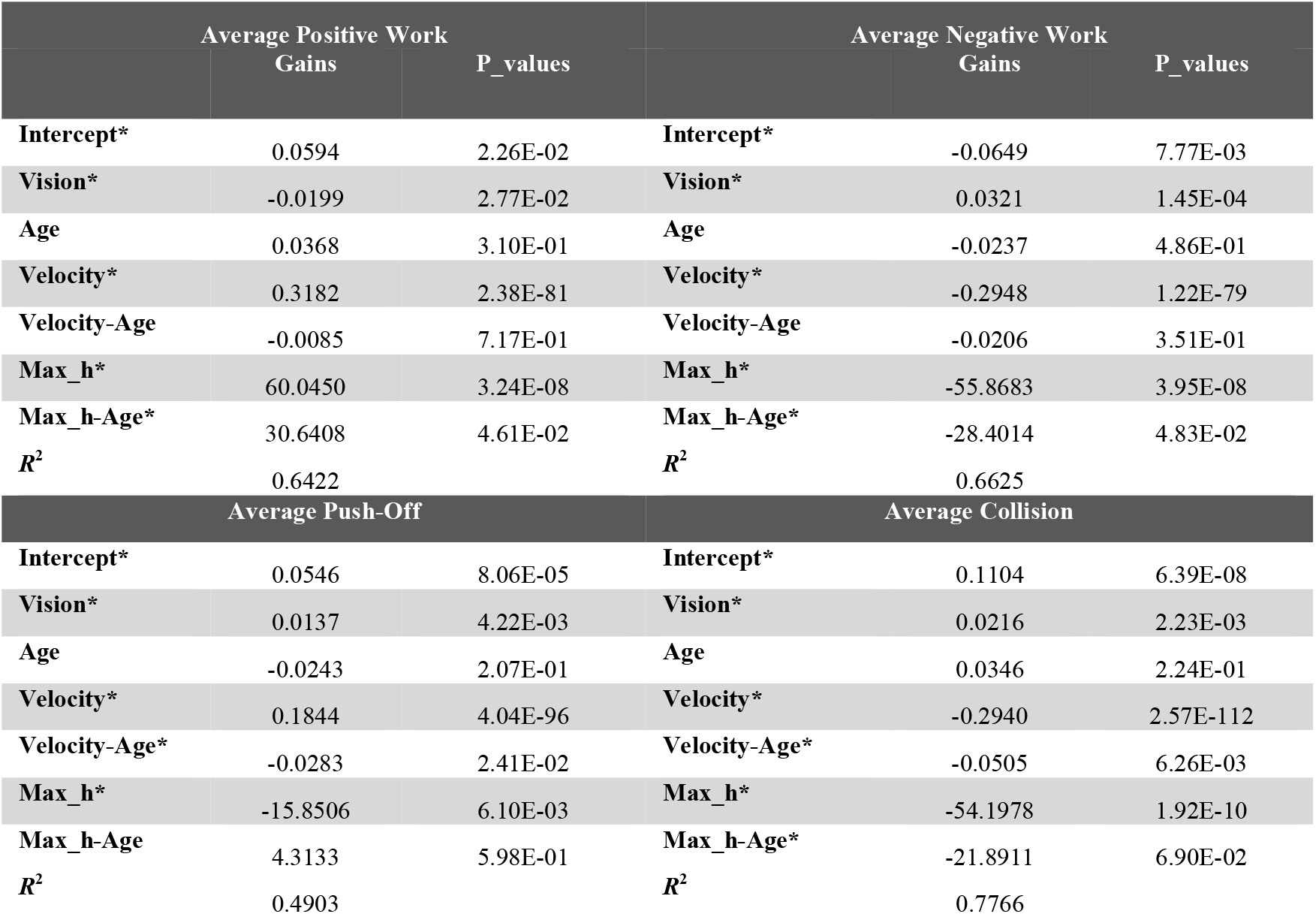
The proposed COM work component trajectories derived by the linear mixed effect model for the experimental data [11]. The significant parameters are identified by an asterisk (*).

**Figure 5.**
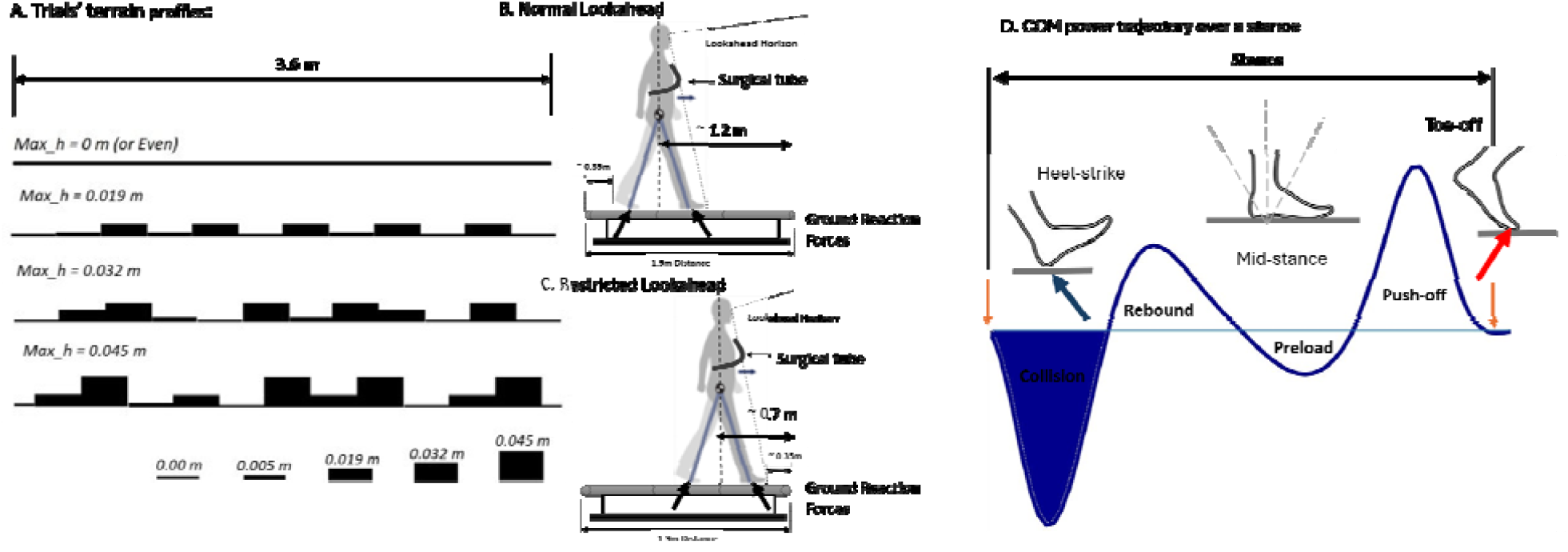
Uneven walking experimental setup [11]: (A) The test terrain profiles for the uneven walking experiment. Each terrain was identified by its maximum foam height. (B) position for the normal lookahead, (C) position for the restricted lookahead. (D) Typical COM power trajectory for a walking stride. It distinctly shows four phases: collision, rebound (single support positive work), preload (single support negative work), and push-off.

We calculated trajectories for total positive work, total negative work, push-off, and collision across walking speeds and terrain amplitudes, ranging from 0.8 m·s□^1^ to 1.6 m·s□^1^ and from 0 m to 0.06 m, respectively.

Trajectories were adjusted to account for random bias in instrumented treadmill calibration. Consequently, we calculated rebound and preload values as follows: rebound as the difference between total positive work and push-off, and preload as the difference between total negative work and collision [10], [11].

Since the trajectories represented various walking conditions where the optimal push-off of each step could vary, we used the “Rebound = Preload” criterion to estimate the preferred walking speed. Additionally, we generated COM work component trajectories for even walking as a validation measure and compared the derived preferred walking speeds (for young and older adults with both normal and restricted lookahead) with values reported in the literature.

## Results

We developed an analytical simulation based on a simple powered walking model [9] to analyze the mechanical energy demand after step-to-step transitions when the push-off does not fully meet the step transition energy requirements, whether for even walking or step ups (uneven walking). The simulation suggested that if push-off does not offset the subsequent collision (in either even or uneven walking) or account for gravitational work (in uneven walking), a mechanical energy deficit will arise post-transition. This deficit must be compensated for during the following step through positive work performance (rebound) [7], [10].

Based on this observation, we inferred that when push-off is suboptimal, the subsequent rebound requires active work, thereby increasing metabolic energy demand. Conversely, if push-off is optimal, the subsequent rebound contributes no extra energy to COM motion, meaning the rebound is passive and may only incur limb oscillation costs [21], [22], [23]. The simulation further indicated that if the push-off impulse exceeds the optimal level, maintaining th prescribed average walking speed and momentum imposes the dissipation of excess mechanical energy post-transition. During the single support phase, this manifests as a larger negative work component (preload) [7], [10] that absorbs the excess energy. When push-off exceeds optimal levels, the rebound is smaller than the preload, suggesting that this excess energy likely converts to heat [24] rather than being stored in limbs and tendons [10], [12]. Thus, excessive push-off results in a rebound that is not entirely passive.

Based on experimentally derived COM work component trajectories, we observed instances where the net single support work dissipated mechanical energy, while in other cases it compensated for mechanical energy deficits. Therefore, for any given walking condition, there was only one walking speed at which rebound and preload were equal. Following simulation results, we assumed that in these cases, no additional positive compensatory work or mechanical dissipation occurred, making the associated single support mechanical work (rebound and preload) passive (storage and release). According to the simulation, we inferred that when the net single support work is zero, the magnitude of mechanical energy dissipation must be minimum. Therefore, we defined the walking speeds under these conditions—taking into account factors such as age, terrain, and visual lookahead—as the preferred walking speeds.

For uneven walking, certain terrain amplitudes within the trial ranges did not yield preferred walking speeds, indicating that the rebound was always greater than the preload (Figure 6). We also reported the range of terrain amplitudes where preferred walking speeds were observed, noting that age and lookahead state decreased the estimated preferred walking speed.

**Figure 6.**
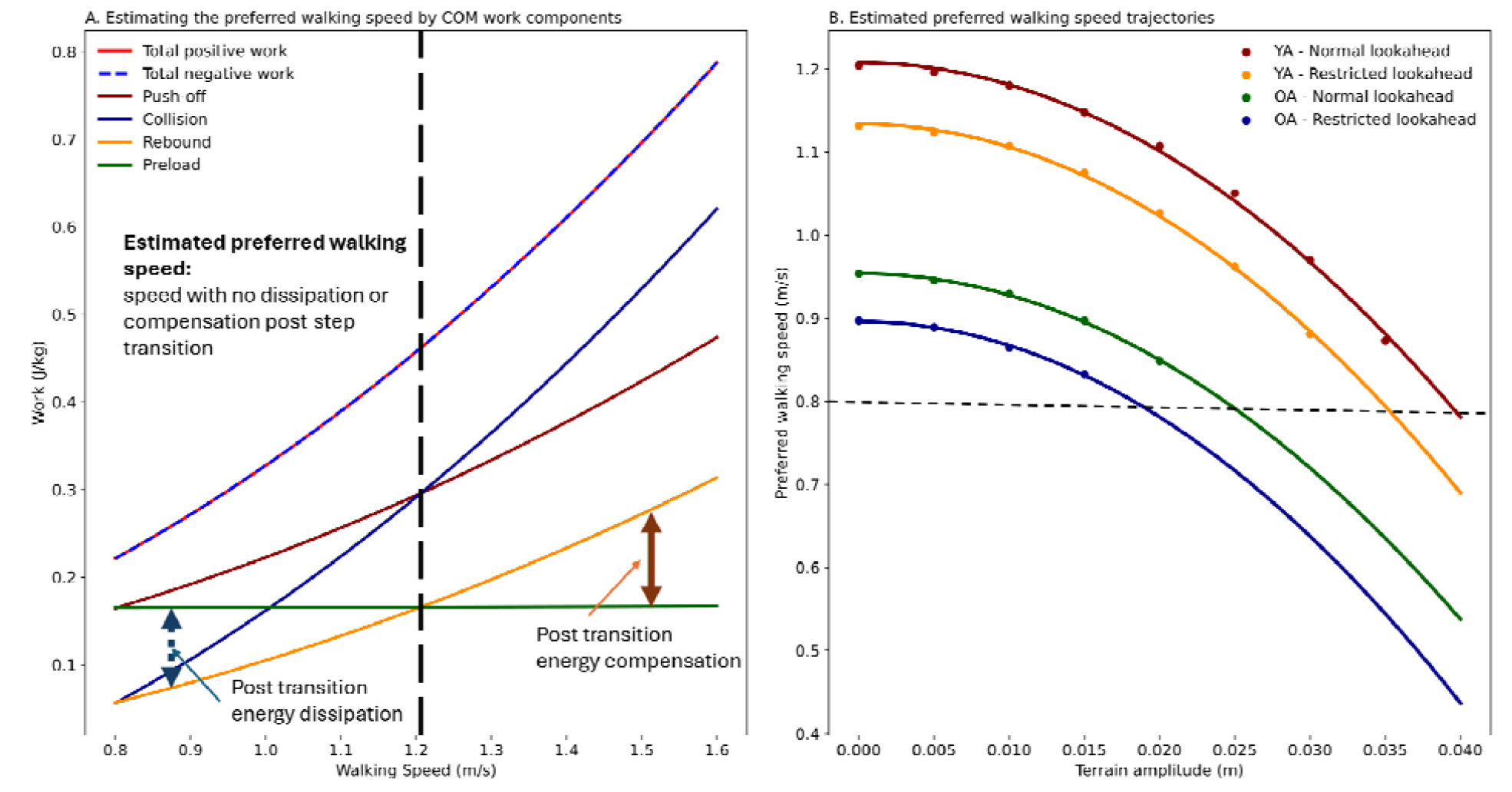
(A) The young adults’ COM work components for even walking with normal lookahead. The crossing points at which the collision or rebound surpasses the push-off or preload is assumed to represent the preferred walking speed at which the entire energy of the step-to-step transition is provided pre-emptively. (B) The estimated preferred walking speed trajectories when terrain complexities increase from zero (even) to 0.045 m. The trajectories indicate terrain amplitude thresholds for each category beyond which the pre-emptive active energy inducement is insufficient to energize the step-to-step transition. Therefore, the rebound work exceeds preload for the entire velocity range (0.8 to 1.6). Some speed estimations are also very low to sustain human walking. Therefore, a lower bound (here 0.8) is set to filter them out.

Second-order polynomials were fitted to the estimated preferred walking speeds within each category as terrain amplitude increased [25]. The preferred walking speed for young adults with normal lookahead, based on “push-off = collision” and “rebound = collision,” was 1.20 m·s□^1^. For young adults with normal lookahead, the preferred walking velocity ranged from 1.20 m·s□^1^ to 0.87 m·s□^1^ as terrain amplitudes rose from 0 m to 0.035 m (−28.1%, p = 4.4e-9, R^2^=0.998).

It is reported that self-selected walking speed declines with visual impairment [26], which we also observed with restricted lookahead. For young adults with restricted lookahead, the preferred walking velocities were estimated to vary from 1.13 m·s□^1^ to 0.88 m·s□^1^ (−22.1%, p = 1.3e-8, R^2^=0.999) as terrain amplitude increased from 0 m to 0.03 m.

The preferred walking speed of older adults is generally lower than that of young adults, which was consistent with our estimates. Additionally, the terrain amplitude ranges for older adults varied based on the state of lookahead. With normal lookahead, terrain amplitudes ranged from 0 m to 0.02 m, and preferred walking speeds declined from 0.95 m·s□^1^ to 0.85 m·s□^1^ (−10.5%, p = 2.2e-5, R^2^=0.999). With restricted lookahead, terrain amplitudes ranged only from 0 m to 0.015 m, and preferred walking speeds declined from 0.9 m·s□^1^ to 0.83 m·s□^1^ (−7.78%, p = 1.7e-3, R^2^=0.997).

## Discussion

For a given walking task or condition, the preferred walking velocity is believed to align with the minimum cost of transport [2]. Optimal walking economy is also suggested to occur when the push-off pre-emptively provides all the energy needed for each step [9]. Since the COM work accounts for most walking energetics [7], the preferred walking speed may also be estimated using COM mechanical work components over the stance phase, mainly by investigating the push-off magnitude. However, finding optimal push-offs for varying walking conditions can be difficult. For example, during uneven walking, where elevation changes frequently, humans must perform work against gravity with varying intensities [27]. Studies indicate that with normal lookahead, humans tend to spread the active work required across multiple steps [28], [29], [30]. Consequently, a specific step’s push-off may exceed the immediate energy need. This requires the exploration of alternatives to directly evaluating push-off work. Here, we consider scenarios where control is exerted based on a “just-in-time” strategy [16].

It is proposed that humans perform both positive and negative work during the single support phase [7], [10]. Given that the single support phase mimics an inverted pendulum’s motion [13], assumed to be conservative, any work performed should theoretically cancel. Otherwise, the COM would either accelerate or decelerate. Hence, in contrast to the pre-emptive push-off, which results from active muscle work, it is proposed that the positive and negative work during the single support phase are passive [7]. In other words, the single support work primarily involves energy storage and release through tendons and tissues [7], [12]. Therefore, if the net single support work is non-zero, it likely serves to regulate COM momentum or energy [11].

Then, identifying speeds where rebound work is fully offset by preload could indicate optimal push-off work for a given walking condition.

Our simulations of step transition impulses and associated work suggest an optimal push-off impulse for each walking speed. This impulse fully supplies the energy required for step transitions, minimizes the total collision dissipation, and eliminates the need for further energy adjustments during the subsequent step. For a given speed, if push-off is suboptimal, additional positive mechanical energy is required; if excessive, energy dissipation must occur post-transition. The COM work component trajectories indicate that under varying conditions (e.g., terrain amplitude, state of lookahead), single support work is not balanced. Hence, this imbalance necessitates net single support mechanical work to regulate COM energy. Consequently, it is plausible to infer that non-zero net single support work arises from increased dissipation either before or after the transition, signaling a rise in metabolic cost. For each walking condition, there is only one speed at which net single support work is zero (rebound = preload). This indicates a push-off that fully offsets heel-strike and gravitational work, resulting in minimal associated dissipation. At these speeds, our simulation suggests that no adjustment to the COM speed is needed post-transition. As soft tissue dissipation during single support (during the rebound and preload phases) is minimal [14], it implies that single support work is passive and incurs no metabolic cost, aside from the potential cost of limb swing due to the frequency of force generation [21], [22], [23].

The estimated preferred walking speeds (self-selected) for even terrain align with reported values for young and older adults [2], [4], [31]. Similar to previous findings [2], preferred speeds for uneven walking are slower than even walking. The estimated speed trends indicate terrain amplitude impact aligns with the quadratic increase in walking costs due to step elevation variations [25], [32]. Moreover, thresholds emerge where no preferred speed exists to pre-emptively meet energy demands. This suggests a biomechanical limitation of the ankle, or interrupted ankle work during uneven walking, making hip torque an additional energy source in such cases. Another observation is the impact of lookahead on even walking, where visual cues allow individuals to optimize footfall choices [16] and synchronize push-off with collisions for energy-efficient steps, or convert maximum potential energy [15], [28], [29]. The importance of the visual information of the coming terrains is evident by slower walking in visually impaired individuals [26], [33], [34]. Some projected speeds are impractically low, suggesting that other walking costs at such speeds might outweigh the benefit of fully energizing gait through push-off [35]. Therefore, low-speed projections, such as those below 0.8 m·s □^1^, should be excluded.

Our method for estimating preferred speeds is based on COM mechanics when the pre-emptive push-off is optimum leading to passive single support work. It aligns with minimizing the COM total dissipative work of each step that closely represents the walking energetics [4]. In our estimation we have considered the terrain amplitude as a varying continuous parameter. However, other factors like time [36], stability, and reproducibility could refine this criterion [37]. In conclusion, although the simple powered model employed cannot directly determine a preferred walking speed, it can offer valuable insights into whether a particular walking regime is mechanically optimal. Since our approach estimates preferred speeds based on COM work trajectories derived from uneven walking experimental data [11], it may potentially be extended to other conditions [7]. In extreme cases when very fast or slow speeds are estimated, biomechanical factors should be examined to ensure walking energetics remain optimal. If no preferred speed is identified, it likely indicates that additional energy regulation is required post-transition, suggesting that push-off alone does not fully energize the gait.

## Notes

Conflict of Interest: None

### Competing Interest Statement

The authors have declared no competing interest.

### Summary of Updates

Modified the analysis/simulation, results, discussion and abstract

